# Pyvolve: A Flexible Python Module for Simulating Sequences along Phylogenies

**DOI:** 10.1101/020214

**Authors:** Stephanie J. Spielman, Claus O. Wilke

## Abstract

We introduce Pyvolve, a flexible Python module for simulating genetic data along a phylogeny using continuous-time Markov models of sequence evolution. Easily incorporated into Python bioinformatics pipelines, Pyvolve can simulate sequences according to most standard models of nucleotide, amino-acid, and codon sequence evolution. All model parameters are fully customizable. Users can additionally specify custom evolutionary models, with custom rate matrices and/or states to evolve. This flexibility makes Pyvolve a convenient framework not only for simulating sequences under a wide variety of conditions, but also for developing and testing new evolutionary models. Pyvolve is an open-source project under a FreeBSD license, and it is available for download, along with a detailed user-manual and example scripts, from http://github.com/sjspielman/pyvolve.

## Introduction

The Python programming language has become a staple in biological computing. In particular, the molecular evolution community has widely embraced Python as standard tool, in part due to the development of powerful bioinformatics modules such as Biopython [1] and DendroPy [2]. However, Python lacks a robust platform for evolutionary sequence simulation.

In computational molecular evolution and phylogenetics, sequence simulation represents a fundamental aspect of model development and testing. Through simulating genetic data according to a particular evolutionary model, one can rigorously test hypotheses about the model, examine the utility of analytical methods or tools in a controlled setting, and assess the interactions of different biological processes [3].

To this end, we introduce Pyvolve, a sequence simulation Python module (with dependencies Biopython [1], SciPy, and NumPy [4]). Pyvolve simulates sequences along a phylogeny using continuous-time Markov models of sequence evolution for nucleotides, amino acids, and codons, according to standard approaches [5]. The primary purpose of Pyvolve is to provide a user-friendly and flexible sequence simulation platform that can easily be integrated into Python bioinformatics pipelines without necessitating the use of third-party software. Furthermore, Pyvolve allows users to specify and evolve custom evolutionary models and/or states, making Pyvolve an ideal engine for novel model development and testing.

## Substitution models and frameworks in Pyvolve

Pyvolve is specifically intended to simulate gene sequences along phylogenies according to Markov models of sequence evolution. Therefore, Pyvolve requires users to provide a fixed phylogeny along which sequences will evolve. Modeling frameworks which are included in Pyvolve are detailed in Table 1.

**Table 1.** Substitution models included in Pyvolve.

| Modeling Framework | Available Models |
| --- | --- |
| Nucleotide | GTR [20] and all nested variants (e.g. HKY85 [21] and TN93 [22]) |
| Amino acid | JTT [23], WAG [24], LG [25], mtMAM [26], mtREV24 [27], DAYHOFF [28], AB [29] |
| Mechanistic codon | GY-style [30,31] and MG-style [32] |
| Empirical codon | ECM (restricted and unrestricted) [33] |
| Mutation-selection | Halpern-Bruno model [10], for both nucleotides and codons |

Pyvolve supports both site-wise and branch (temporal) heterogeneity. Site-wise heterogeneity can be modeled with Γ or Γ+I rates, or users can specify a custom rate-distribution. Further, users can specify a custom rate matrix for a given simulation, and thus they can evolve sequences according to substitution models other than those shown in Table 1. Similarly, users have the option to specify a custom set of states to evolve, rather than being limited to nucleotide, amino-acid, or codon data. Therefore, it is possible to specify arbitrary models with corresponding custom states, e.g. states 0, 1, and 2. This general framework will enable users to evolve, for instance, states according to models of character evolution, such as the Mk model [6].

Similar to other simulation platforms (e.g. Seq-Gen [7], indel-Seq-Gen [8], and INDELible [9]), Pyvolve simulates sequences in groups of partitions, such that different partitions can have unique evolutionary models and/or parameters. Although Pyvolve enforces that all partitions within a single simulation evolve according to the same model family (e.g. nucleotide, amino-acid, or codon), Python’s flexible scripting environment allows for straight-forward alignment concatenation. Therefore, it is readily possible to embed a series of Pyvolve simulations within a Python script to produce highly-heterogeneous alignments, for instance where coding sequences are interspersed with non-coding DNA sequences. Moreover, Pyvolve allows users to specify, for a given partition, the ancestral sequence (MRCA) to evolve.

In addition, we highlight that Pyvolve is among the first open-source simulation tools to include the mutation-selection modeling framework introduced by Halpern and Bruno in ref. [10] (we note that the simulation software SGWE [11] also includes this model). Importantly, although these models were originally developed for codon evolution [10, 12], Pyvolve implements mutation-selection models for both codons and nucleotides. We expect that this implementation will foster the continued development and use of this modeling framework, which has seen a surge of popularity in recent years [13–19].

## Basic Usage of Pyvolve

The basic framework for a simple simulation with the Pyvolve module is shown in Fig. 1. To simulate sequences, users should input the phylogeny along which sequences will evolve, define evolutionary model(s), and assign model(s) to partition(s). Pyvolve implements all evolutionary models in their most general forms, such that any parameter in the model may be customized. This behavior stands in contrast to several other simulation platforms of comparable scope to Pyvolve. For example, some of the most commonly used simulation tools that implement codon models, including INDELible [9], EVOLVER [34], and PhyloSim [35], do not allow users to specify dS rate variation in codon models. Pyvolve provides this option, among many others.

**Figure 1.**
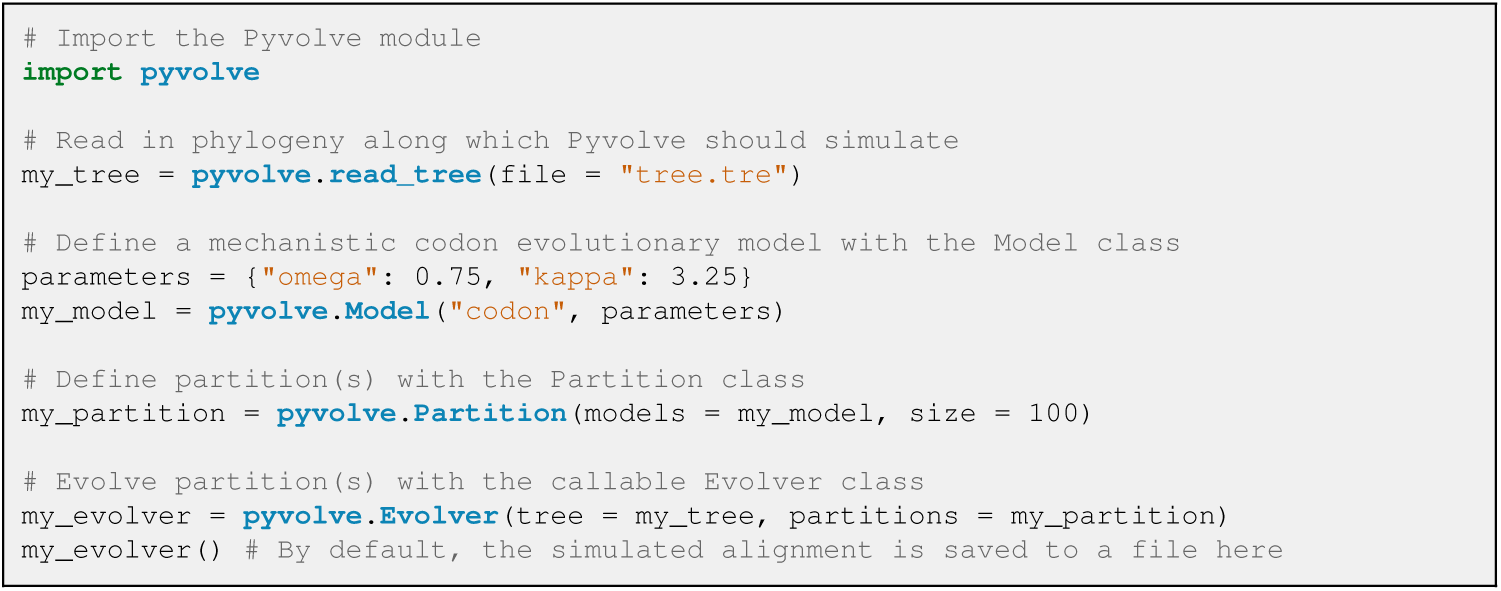
**Example code for a simple codon simulation in Pyvolve**. This example will simulate an alignment of 100 codons with a *dN/dS* of 0.75 and a *?* (transition-tranversion mutational bias) of 3.25. By default, sequences will be output to a file called “simulated alignment.fasta”, although this file name can be changed, as described in Pyvolve’s user manual.

In the example shown in Fig. 1, stationary frequencies are not explicitly specified. Under this circumstance, Pyvolve will assume equal frequencies, although they would normally be provided using the key 

~~~
“state_freqs”
~~~

 in the dictionary of parameters. Furthermore, Pyvolve contains a convenient module to help specify state frequencies. This module can read in frequencies from an existing sequence and/or alignment file (either globally or from specified alignment columns), generate random frequencies, or constrain frequencies to a given list of allowed states. In addition, this module will convert frequencies between alphabets, which is useful, for example, if one wishes to simulate amino-acid data using the state frequencies as read from a file of codon sequence data.

## Validating Pyvolve

We carefully assessed that Pyvolve accurately simulates sequences. To this end, we simulated several data sets under a variety of evolutionary models and conditions and compared the observed substitution rates with the simulated parameters.

To evaluate Pyvolve under the most basic of conditions, site-homogeneity, we simulated both nucleotide and codon data sets. We evolved nucleotide sequences under the JC69 model [36] across several phylogenies with varying branch lengths (representing the substitution rate), and we evolved codon sequences under a MG94-style model [32] with varying values of *dN/dS*. All alignments were simulated along a two-taxon tree and contained 100,000 positions. We simulated 50 replicates for each branch length and/or *dN/dS* parameterization. As shown in Figs. 2A and B, the observed number of changes agreed precisely with the specified parameters.

**Figure 2.**
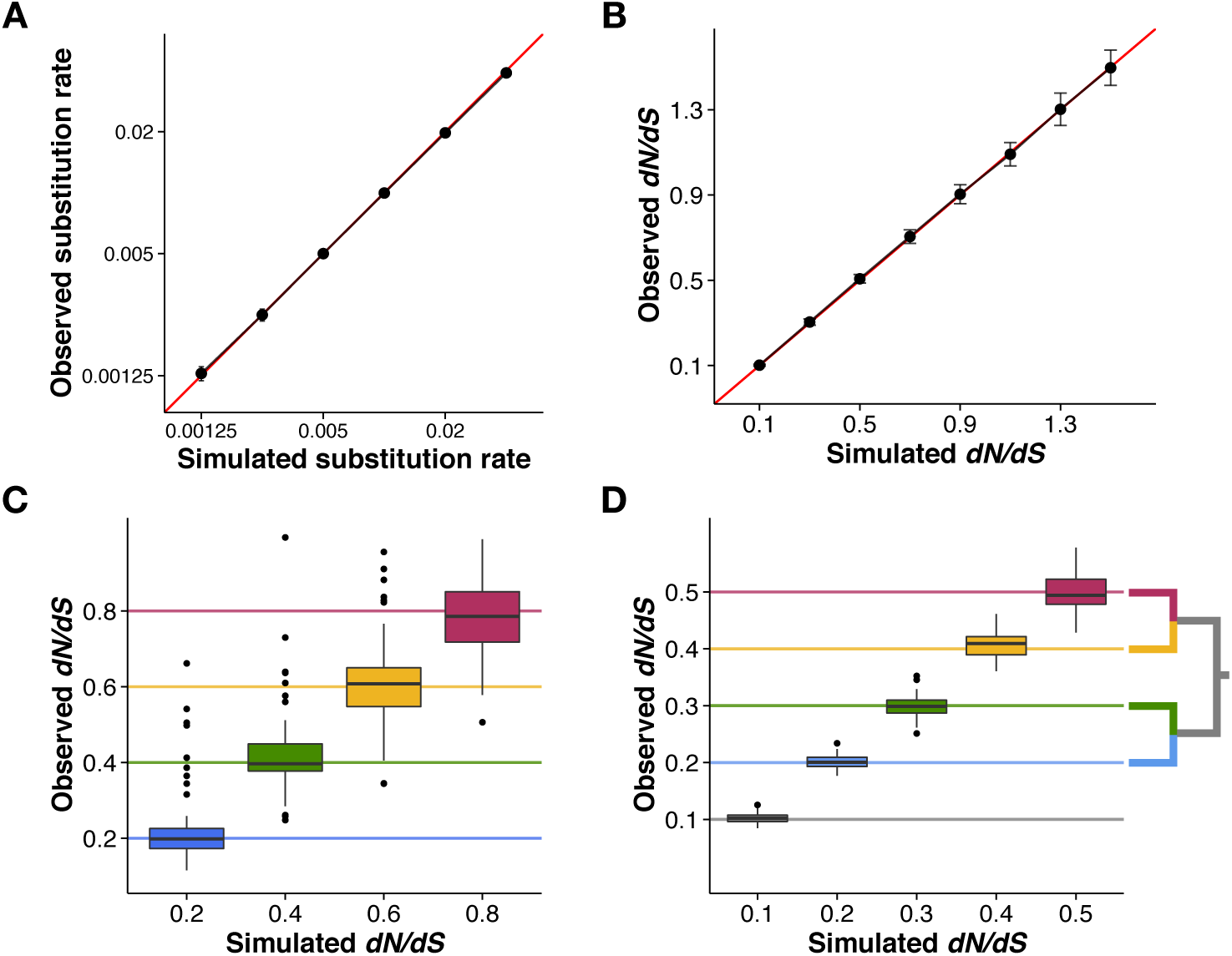
**Pyvolve accurately evolves sequences under homogenous, site-wise rate heterogeneity, and branch-specific rate heterogeneity**. A) Nucleotide alignments simulated under the JC69 [36] model along two-taxon trees with varying branch lengths, which represent the substitution rate. Points represent the mean observed substitution rate for the 50 alignment replicates simulated under the given value, and error bars represent standard deviations. The red line indicates the *x = y* line. B) Codon alignments simulated under an MG94-style [32] model with varying values for the *dN/dS* parameter. Points represent the mean *dN/dS* inferred from the 50 alignment replicates simulated under the given *dN/dS* value, and error bars represent standard deviations. The red line indicates the *x = y* line. C) Site-wise heterogeneity simulated with an MG94-style [32] model with varying *dN/dS* values across sites. Horizontal lines indicate the simulated *dN/dS* value for each *dN/dS* category. D) Branch-wise heterogeneity simulated with an MG94-style [32] model with each branch evolving according to a distinct *dN/dS* value. Horizontal lines indicate the simulated *dN/dS* value for each branch, as shown in the inset phylogeny. The lowest *dN/dS* category (*dN/dS* =0.1) was applied to the internal branch (shown in gray). All code and data used to validate Pyvolve’s performance and generate this figure are available in File S1.

We additionally validated Pyvolve’s implementation of site-wise rate heterogeneity. We simulated an alignment of 400 codon positions, again under an MG94-style model [32], along a balanced tree of 2^14^ taxa with all branch lengths set to 0.01. This large number of taxa was necessary to achieve accurate estimates for site-specific measurements. To incorporate site-specific rate heterogeneity, we specified four *dN/dS* values of 0.2, 0.4, 0.6, and 0.8, to be assigned in equal proportions to sites across this alignment. We counted the observed *dN/dS* values for each resulting alignment column using a version of the Suzuki-Gojobori algorithm [37] adapted for phylogenetic data [38]. Figure 2C demonstrates that Pyvolve accurately implements site-specific rate heterogeneity. The high variance seen in Figure 2C is an expected result of enumerating substitutions on a site-specific basis, which, as a relatively small data set, produces substantial noise.

Finally, we confirmed that Pyvolve accurately simulates branch heterogeneity. Using a four-taxon tree, we evolved codon sequences under an MG94-style model [32] and specified a distinct *dN/dS* ratio for each branch. We simulated 50 replicate alignments of 100,000 positions, and we computed the observed *dN/dS* value along each branch. Figure 2D shows that observed branch *dN/dS* values agreed with the simulated values.

## Conclusions

Because Pyvolve focuses on simulating the substitution processes using continuous-time Markov models along a fixed phylogeny, it is most suitable for simulating gene sequences, benchmarking inference frameworks, and for developing and testing novel Markov models of sequence evolution. For example, we see a primary application of Pyvolve as a convenient simulation platform to benchmark *dN/dS* and mutation-selection model inference frameworks such as the ones provided by PAML [34], HyPhy [39], Phylobayes [17], or swMutSel [16]. Indeed, the Pyvolve engine has already successfully been applied to investigate the relationship between mutation-selection and *dN/dS* modeling frameworks and to identify estimation biases in certain *dN/dS* models [18]. Moreover, we believe that Pyvolve provides a convenient tool for easy incorporation of complex simulations, for instance those used in approximate Bayesian computation (ABC) or MCMC methods [40], into Python pipelines.

Importantly, Pyvolve is meant primarily as a convenient Python library for simulating simple Markov models of sequence evolution. For more complex evolutionary scenarios, including the simulation of entire genomes, population processes, or protein folding and energetics, we refer the reader to several more suitable platforms. For example, genomic process such as recombination, coalescent-based models, gene duplication, and migration, may be best simulated with softwares such as ALF [41], CoalEvol and SGWE [11], or EvolSimulator [42]. Simulators which consider the influence of structural and/or biophysical constraints in protein sequence evolution include CASS [43] or ProteinEvolver [44]. Similarly, the software REvolver [45] simulates protein sequences with structural domain constraints by recruiting profile hidden Markov models (pHMMs) to model site-specific substitution processes.

We additionally note that Pyvolve does not currently include the simulation of insertions and deletions (indels), although this functionality is planned for a future release. We refer readers to the softwares indel-Seq-Gen [8] and Dawg [46] for simulating nucleotide sequences, and we suggest platforms such as INDELible [9], phyloSim [35], or πBuss [47] for simulating coding sequences with indels.

In sum, we believe that Pyvolve’s flexible platform and user-friendly interface will provide a helpful and convenient tool for the biocomputing Python community. Pyvolve is freely available from http://github.com/sjspielman/pyvolve, conveniently packaged with a comprehensive user manual and several example scripts demonstrating various simulation conditions. In addition, Pyvolve is distributed with two helpful Python scripts that complement Pyvolve simulations: one which implements the Suzuki-Gojobori [37] *dN/dS* counting algorithm adapted for phylogenetic data [38], and one which calculates *dN/dS* from a given set of mutation-selection model parameters as described in ref. [18]. Pyvolve is additionally available for download from the Python Package Index (e.g. via pip).

## Supporting Information

### File S1

**This file contains all scripts and data used to validate Pyvolve.**

## Acknowledgments

We thank Suyang Wan and Dariya Sydykova for helpful feedback during Pyvolve development and Rebecca Tarvin for her help designing the Pyvolve logo.

